# GIP receptor agonism suppresses inflammation-induced aversion and food intake via distinct circuits

**DOI:** 10.1101/2025.08.12.669936

**Authors:** Haley S. Province, Nikolas W. Hayes, Nathan A. Leong, Carolyn M. Lorch, Alexandra Pekerman, Jessica L. Xia, Lisa R. Beutler

## Abstract

Glucose-dependent insulinotropic polypeptide (GIP) is a gut-derived incretin hormone, and pharmacologic modulation of central GIP receptors (GIPR) improves energy homeostasis. Recent reports have demonstrated that GIPR agonism is also anti-aversive. However, the mechanisms by which GIPR signaling impact food intake and aversion are incompletely understood. Here, we show that GIPR agonism abrogates the aversive and enhances the anorexigenic effects of the pro-inflammatory cytokine interleukin-1β (IL-1β). Aversion-encoding parabrachial calcitonin-gene related peptide (CGRP) neurons were required for IL-1β-induced conditioned taste avoidance (CTA) but not anorexia. Moreover, systemic IL-1β increased *in vivo* CGRP neural activity, and this was significantly attenuated by co-administration of a GIPR agonist. By contrast, GIPR in the dorsal vagal complex were required for the acute anorectic effect of GIPR agonism but not its anti-aversive effect. Taken together, our data suggest that GIPR agonism reduces food intake and prevents aversion via distinct circuits, and that GIPR agonism may represent an effective approach to alleviate inflammation-induced aversion.

## INTRODUCTION

Glucose-dependent insulinotropic polypeptide (GIP) is a gut-derived incretin hormone that acts on GIP receptors (GIPR) to regulate energy homeostasis and glucose-dependent insulin secretion. GIPR are expressed peripherally in pancreatic islets and adipose tissue and centrally in hypothalamic feeding nuclei and the dorsal vagal complex (DVC) [1–4]. Pharmacologic manipulation of central GIPR signaling alters energy homeostasis, with both agonism and antagonism of GIPR decreasing body weight and food intake [5–10]. Recent reports have also demonstrated a novel role for pharmacologic GIPR agonism in preventing conditioned taste avoidance (CTA) caused by satiety hormones, lithium chloride, and growth and differentiation factor-15 (GDF-15) [11–13]. Additionally, GIPR agonism significantly reduces emesis due to liraglutide and cisplatin in preclinical models and decreases liraglutide-induced gastrointestinal adverse events in humans, suggesting a potential role for GIPR agonists as anti-aversive agents in the clinic [14–17]. However, the extent to which GIPR agonism modulates inflammation-induced aversion is unknown.

Inflammation causes non-specific and unpleasant sickness behaviors, including anorexia, lethargy and fever. The intrinsically aversive nature of the inflammatory response is evolutionarily advantageous, facilitating behavioral adaptations to avoid the source of illness [18–20]. However, for those experiencing chronic inflammation, this can become maladaptive and contribute to reduced quality of life [21, 22]. Recent studies have identified some of the neural circuit mechanisms that drive the behavioral changes caused by peripheral inflammation [23–26], but these processes remain incompletely understood. Further, there is an unmet need for effective anti-aversive pharmacotherapies to alleviate symptoms associated with acute and chronic inflammation.

Interleukin-1β (IL-1β) is a pro-inflammatory cytokine that is elevated in infections and autoimmune diseases [19, 27] and induces robust sickness behavior in mice [28, 29]. To determine the impact of GIPR agonism on inflammation-induced behavioral changes and neuron activity, we peripherally administered IL-1β to acutely induce an aversive inflammatory state. GIPR agonism prevented IL-1β-induced CTA. In agreement, GIPR agonism reduced *in vivo* IL-1β-induced activation of aversion-promoting calcitonin-gene related peptide-expressing neurons (CGRP neurons) in the parabrachial nucleus [30–33]. By contrast, GIPR agonism modestly potentiated IL-1β-induced anorexia via direct action on GIPR in the DVC, which were dispensable for GIP-induced anti-aversion. Combined, our findings illuminate the neural circuit basis underlying multiple behavioral effects of GIPR agonism and reveal a potential role for GIPR agonism in the management of inflammation-induced aversive symptoms.

## RESULTS

### IL-1β activates CGRP neurons, and this is required for IL-1β-induced CTA

Inflammatory stimuli, such as lipopolysaccharide (LPS), tumor necrosis factor-α (TNF-α), and interleukin-1β (IL-1β) induce an array of aversive sickness behaviors such as anorexia, malaise and lethargy [28, 29]. While the central pathways underlying the aversive effects of inflammation are not fully understood, many types of aversive stimuli activate parabrachial CGRP neurons. Peripheral administration of LPS, TNF-α, and IL-1β each induce the expression of Fos, a common marker of neural activation, in parabrachial CGRP neurons [34]. In addition, these aversion-promoting neurons are critical for establishing CTA caused by LPS and GDF-15 [31–33, 35]. To understand how inflammatory stimuli impact *in vivo* CGRP neuron activity, we used fiber photometry to record CGRP neuron calcium activity (**Fig. S1A–B**). Intraperitoneal (i.p.) administration of LPS, TNF-α, and IL-1β each increased CGRP neuron calcium activity compared to phosphate-buffered saline (PBS) vehicle injection, an effect that was most pronounced for IL-1β (**Fig. 1A–L**). This IL-1β-induced increase in CGRP neuron activity was equally robust upon a second administration of IL-1β after a washout period (**Fig. S1C–F**).

**Figure 1.**
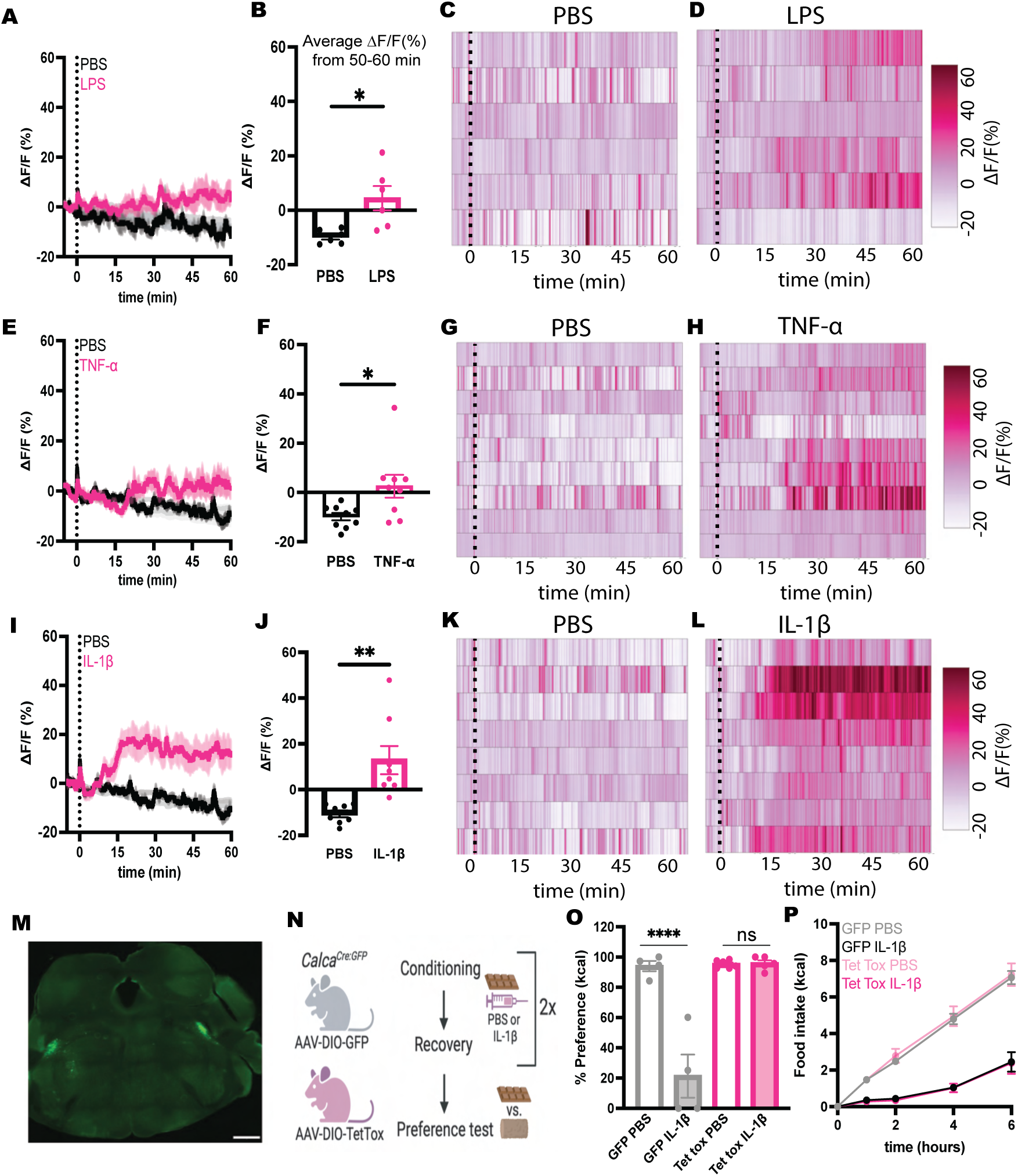
IL-1β activates CGRP neurons, and this is required for IL-1β-induced CTA. **(A, E, I)** Calcium signal in CGRP neurons from fasted mice injected with PBS vehicle control (black traces) or LPS (500 ug/kg, **A**), TNF-α (200 ug/kg, **E**), or IL-1β (10 ug/kg, **I**). N = 6-9 mice per group. For all recordings, isosbestic traces are shown in gray, vertical dashed lines indicate the time of injection, and traces indicate mean ± SEM. **(B, F, J)** Average ΔF/F (%) 50-60 min after injection in mice from **(A, E, I)**. Paired t-tests: **B**: p=0.0238; **F:** p = 0.0298; **J:** p = 0.0068. **(C, D, G, H, K, L)** Heat maps showing ΔF/F in individual mice injected with PBS, LPS, TNF-α, or IL-1β as indicated. Each heatmap line represents an individual animal. **(M)** Representative image of TetTox-GFP expression in parabrachial CGRP neurons in a *Calca^Cre:GFP^* mouse. Scalebar is 1 mm. **(N)** Schematic of CTA experiment. **(O)** Percent preference for chocolate versus standard chow in GFP and TetTox-injected *Calca^Cre:GFP^*mice conditioned with PBS or IL-1β (5 μg /kg) i.p. as indicated. N = 4-6 mice per group. Two-way ANOVA drug x virus interaction effect p<0.0001. Post-hoc comparisons: ****p<0.0001. **(P)** Food intake in fasted GFP and TetTox-injected *Calca^Cre:GFP^* mice following acute PBS or IL-1β (5 μg /kg) i.p. N = 8-10 mice per group. Two-way ANOVA of total kcal drug x virus interaction effect p=0.8354. **(A, B, E, F, I, J, O, P)** Data are shown as mean ± SEM.

As expected, IL-1β induced robust CTA (**Fig. 1O, Fig. S1G**) [36, 37]. To determine whether CGRP neuron activity is required for IL-1β-induced CTA, we injected a Cre-dependent viral vector encoding tetanus toxin light chain (Tet Tox) bilaterally into the lateral parabrachial nucleus of mice that express Cre recombinase in CGRP neurons, permanently blocking neurotransmitter release from these neurons (**Fig. 1M**). Tet Tox and GFP control mice underwent two conditioning sessions where they consumed a novel and palatable food (chocolate), followed by IL-1β or PBS control injection (**Fig. 1N**). After a two-day drug washout period, mice underwent preference testing where they had access to both chocolate and standard chow for 30 minutes without further drug administration. While control mice developed a robust CTA when chocolate was paired with IL-1β, mice with silenced CGRP neurons did not (**Fig. 1O**). We observed a similar effect using the non-caloric sweetener saccharin instead of chocolate (**Fig. S1G**). Surprisingly, the acute anorexic effect of IL-1β treatment was indistinguishable between Tet Tox and GFP control mice (**Fig. 1P**). In summary, CGRP neurons are activated by inflammatory stimuli and are required for IL-1β-induced CTA, but not anorexia.

### GIPR agonism decreases IL-1β-induced CTA and CGRP neuron activation

GIPR agonism has been shown to attenuate CTA to a variety of aversive stimuli [12–15]. Further, GIPR agonism decreases Fos expression in the parabrachial nucleus induced by peripheral administration of the satiety hormone peptide-YY [12]. However, the mechanism by which GIPR agonism prevents aversion during inflammation is not fully understood. Therefore, we next determined whether GIPR agonism modulates IL-1β-induced aversive behaviors and parabrachial CGRP neuron activity. C57BL/6 mice developed CTA to chocolate when paired with IL-1β. Co-administration of the GIPR agonist D-Ala^2^-GIP (DA-GIP) during conditioning significantly attenuated IL-1β-induced CTA (**Fig. 2A**). To test whether GIPR agonism alters CGRP neuron activity, we used fiber photometry to record calcium activity in CGRP neurons in mice following PBS or DA-GIP injection. DA-GIP alone had no effect on CGRP neuron activity in either fasted or fed mice (**Fig. 2B–D, Fig. S2A–C**). However, co-administration of DA-GIP and IL-1β significantly attenuated IL-1β-induced CGRP neuron activation throughout the duration of the recording (**Fig. 2E–H**). As DA-GIP reduced both CTA and CGRP neuron response to IL-1β, we conclude that GIPR agonism modulates both the behavioral and neural responses to inflammation.

**Figure 2.**
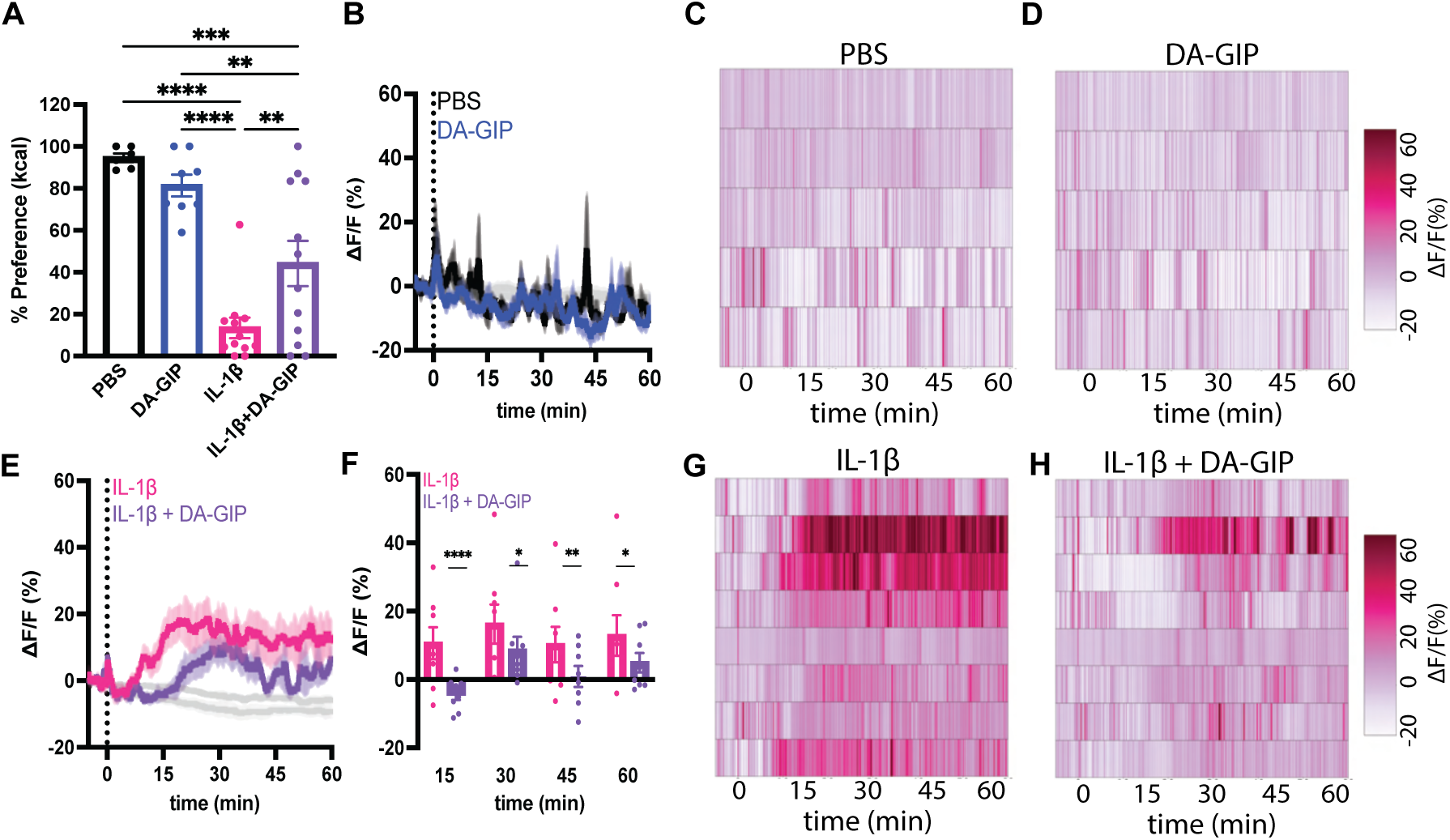
GIP receptor agonism decreases IL-1β-induced conditioned taste avoidance and CGRP neuron activation. **(A)** Percent preference for chocolate versus standard chow in C57BL/6 mice conditioned with PBS, DA-GIP (1 mg/kg), IL-1β (5 μg /kg), or DA-GIP + IL-1β i.p. as indicated. N = 6-12 mice per group. One-way ANOVA p< 0.0001. **(B)** Calcium signal in CGRP neurons from fasted mice injected with PBS or DA-GIP (1 mg/kg) i.p. N = 5 mice per group. **(C–D)** Heat maps showing ΔF/F in individual mice injected with PBS or DA-GIP. **(E)** Calcium signal in CGRP neurons from fasted mice injected with IL-1β (10 µg/kg) or IL-1β + DA-GIP i.p. N = 8 mice per group. **(F)** Average ΔF/F in mice from **(E)** 15, 30, 45, and 60 minutes after injection. Two-way ANOVA effect of drug p=0.0317. **(G–H)** Heat maps showing ΔF/F in individual mice injected with IL-1β or DA-GIP + IL-1β. **(A, F)** Data are shown as mean ± SEM. Post-hoc comparisons: *p<0.05, **p < 0.01; ***p<0.001, ****p<0.0001. **(B, E)** Isosbestic traces for all recordings are shown in gray. Vertical dashed lines indicate the time of injection. Traces indicate mean ± SEM. **(C, D, G, H)** Each heatmap line represents an individual animal.

### GIPR agonism and 5-HT_3_ receptor antagonism additively suppress IL-1β-induced CTA and CGRP neuron activation

Currently, GIPR agonists are not used clinically to minimize aversive symptoms. One mainstay treatment for aversive symptoms is the 5-HT_3_ receptor antagonist ondansetron (OND), which can alleviate emesis and nausea [38]. The ability of OND to prevent CTA in rodent models is controversial, with studies reporting that OND does not attenuate CTA due to serotonin, cisplatin and ipecac [39, 40] and another reporting that OND prevents CTA due to apomorphine [41]. To determine whether the anti-aversive properties of DA-GIP and OND have overlapping neural substrates, we next evaluated the effects of OND. OND alone did not attenuate IL-1β-induced CTA (**Fig. S3A**), but significantly reduced IL-1β-driven CGRP neuron activation, despite having no effect on CGRP neuron activity when administered alone (**Fig. S3B–H**, **Fig. S2D–F**). However, the CGRP neuron activation by IL-1β in the presence or absence of OND co-treatment was not significantly different by one-hour after injection (**Fig. S3F**). This may explain the inability of ondansetron to attenuate CTA, as animals can develop a CTA with up to an hour delay between the taste and the onset of sickness [42, 43]. Combined DA-GIP and OND treatment almost fully attenuated IL-1β-induced CTA and CGRP neuron activation (**Fig. 3A–E**).

**Figure 3.**
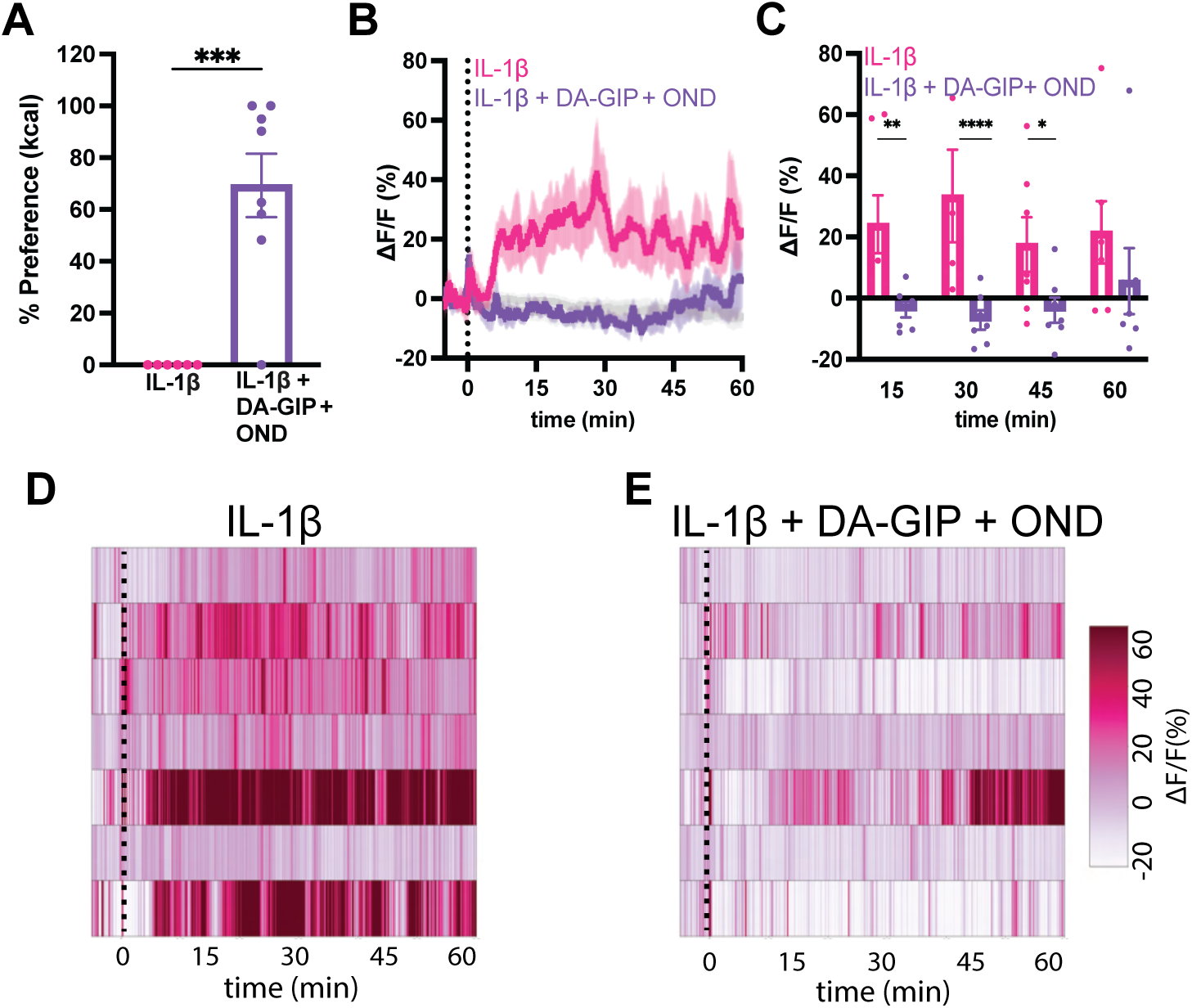
DA-GIP and ondansetron additively suppress IL-1β-induced conditioned taste avoidance and CGRP neuron activation. **(A)** Percent preference for chocolate versus standard chow in C57BL/6 mice conditioned with IL-1β (5 μg /kg) or IL-1β + DA-GIP (1 mg/kg) + ondansetron (OND, 1 mg/kg) i.p. as indicated. N = 6-8 mice per group. Unpaired T test p=0.0004. **(B)** Calcium signal in CGRP neurons from fasted mice injected with IL-1β (10 µg/kg), or IL-1β + ondansetron (OND, 1 mg/kg) + DA-GIP (1 mg/kg) i.p. N = 7 mice per group. **(C)** Average ΔF/F in mice from **(B)** 15, 30, 45, and 60 minutes after injection. Two-way ANOVA effect of drug p=0.0388. Post-hoc comparisons: *p<0.05, **p<0.01. **(D–E)** Heat maps showing ΔF/F in individual mice injected with IL-1β or IL-1β + DA-GIP + OND. Each heatmap line represents an individual animal. **(A, C)** Data are shown as mean ± SEM.

Taken together, these results suggest that DA-GIP and OND exert their anti-aversive effects via distinct mechanisms, and that GIPR agonism may be a useful adjunctive approach to further alleviate aversive inflammation symptoms. Finally, our results argue that the extent to which a drug suppresses CGRP neuron activation may more accurately predict the clinical efficacy of anti-aversive or anti-emetic compounds than CTA in rodent models.

### Anti-inflammatory drug treatment fails to attenuate IL-1β-induced CTA and CGRP neuron activation

We next assessed whether IL-1β-induced behavioral and neural changes could be attenuated by non-steroidal anti-inflammatory drugs (NSAIDs) that work by inhibiting prostaglandin synthesis. NSAIDs have previously been shown to rescue inflammation-induced anorexia but not CTA [44, 45]. Concordant with published results, pre-treatment with the non-selective NSAID ketorolac (KET) did not prevent IL-1β-induced CTA (**Fig. S4A**). Further, KET did not alter CGRP neuron activity, nor did it significantly attenuate IL-1β-induced CGRP neuron activation (**Fig. S4C–H)**. These findings suggest that IL-1β induces CTA and CGRP neuron activity through a prostaglandin-independent mechanism.

### DA-GIP, OND, and KET have distinct effects on IL-1β-induced anorexia

Numerous reports have shown that aversion and food-intake suppression are mediated by separate neural circuits [44, 46–48]. However, whether blocking the development of aversion is necessarily accompanied by reduced anorexia is unclear. To explore this, we tested how DA-GIP influences acute IL-1β-induced anorexia. DA-GIP alone modestly reduced food intake and potentiated anorexia when co-administered with IL-1β, consistent with previous reports that GIPR agonism reduces food intake when paired with other anorexigenic stimuli [11, 12, 49] (**Fig. 4A**). OND had no effect on food intake when given alone or co-administered with IL-1β (**Fig. 4B**). As reported by others, pretreatment with KET restored food intake in mice administered IL-1β (**Fig. 4C**) [44, 45], despite having no effect on CTA or inflammation-induced CGRP neuron activation (**Fig. S4A**). Altogether, these data demonstrate that inflammation-induced aversion and anorexia are dissociable.

**Figure 4.**
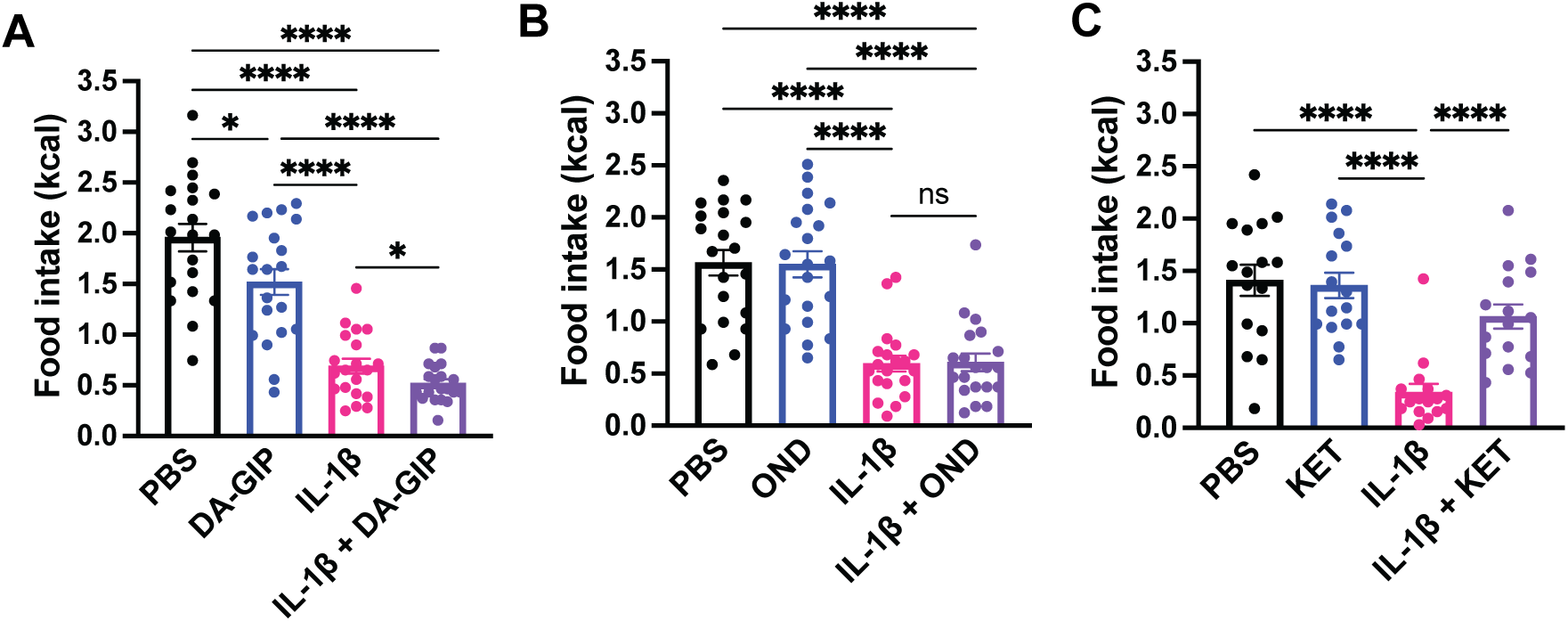
DA-GIP, ondansetron, and ketorolac have distinct effects on IL-1β-induced anorexia. **(A–C)** Food intake in fasted C57BL/6 mice injected with PBS, DA-GIP (1 mg/kg), IL-1β (5 μg /kg), or IL-1β + DA-GIP i.p. **(A)**, PBS, OND (1 mg/kg), IL-1β, or IL-1β + OND i.p. **(B)**, or PBS, ketorolac (KET, 15 mg/kg), IL-1β, or IL-1β + KET i.p. **(C)** as indicated. Food intake commenced at the onset of dark cycle and was measured for 2 hours. One-way repeated measures ANOVA: **A:** p<0.0001; **B**: p<0.0001; **C**: p<0.0001. N = 16-20 mice per group. Post-hoc comparisons: *p<0.05, ****p<0.0001. Data are shown as mean ± SEM.

### GIPR in DVC neurons are required for DA-GIP-induced anorexia but not anti-aversion

We next set out to determine which GIPR-expressing neurons mediate the anti-aversive and/or the anorectic effect DA-GIP. Previous work demonstrated that GIPR are expressed on inhibitory interneurons in the area postrema (AP), a region known to coordinate aversive and emetic responses [50]. GIPR-expressing neurons project within the AP to inhibit the output of aversive neuron populations to other parts of the brain, including parabrachial CGRP neurons, and ablating AP GIPR neurons attenuates the anti-aversive effect of GIPR agonism [13, 35, 48]. By contrast, chemogenetic activation of GIPR neurons in the dorsal vagal complex (DVC), comprised of the AP and nucleus of the solitary tract (NTS), induces CTA and decreases food intake [1]. However, no studies have directly assessed the role of DVC GIPR themselves in GIP-mediated anorexia and anti-aversion.

To test this, we injected AAV-Cre-GFP into the DVC of *Gipr*^lox/lox^ and *Gipr*^+/+^ littermate control mice to selectively delete *Gipr* [51]. Though we initially aimed to selectively target the AP, virus often spread to the surrounding NTS (**Fig. S5**). Animals included in the final analysis had strong GFP expression in the AP with some spillover into the neighboring NTS (**Fig. 5A**, **S5**). DA-GIP treatment induced pronounced Fos expression in the AP of AAV-Cre-GFP-injected *Gipr*^+/+^ mice compared to vehicle injection, and Fos induction was significantly attenuated in *Gipr*^lox/lox^ mice (**Fig. 5H–I**). Notably, the presence of some DA-GIP-induced Fos induction in the AP of *Gipr*^lox/lox^ mice suggests either that *Gipr* was not deleted in all AP neurons or that DA-GIP activates a subset of those neurons indirectly. DA-GIP did not dramatically increase Fos expression in the caudal NTS (measured at the level of the AP) of *Gipr*^+/+^ mice (**Fig. S6**), consistent with a prior report that peripherally administered GIPR agonists preferentially access the circumventricular organs and that GIPR are more robustly expressed in AP than in NTS [1].

**Figure 5.**
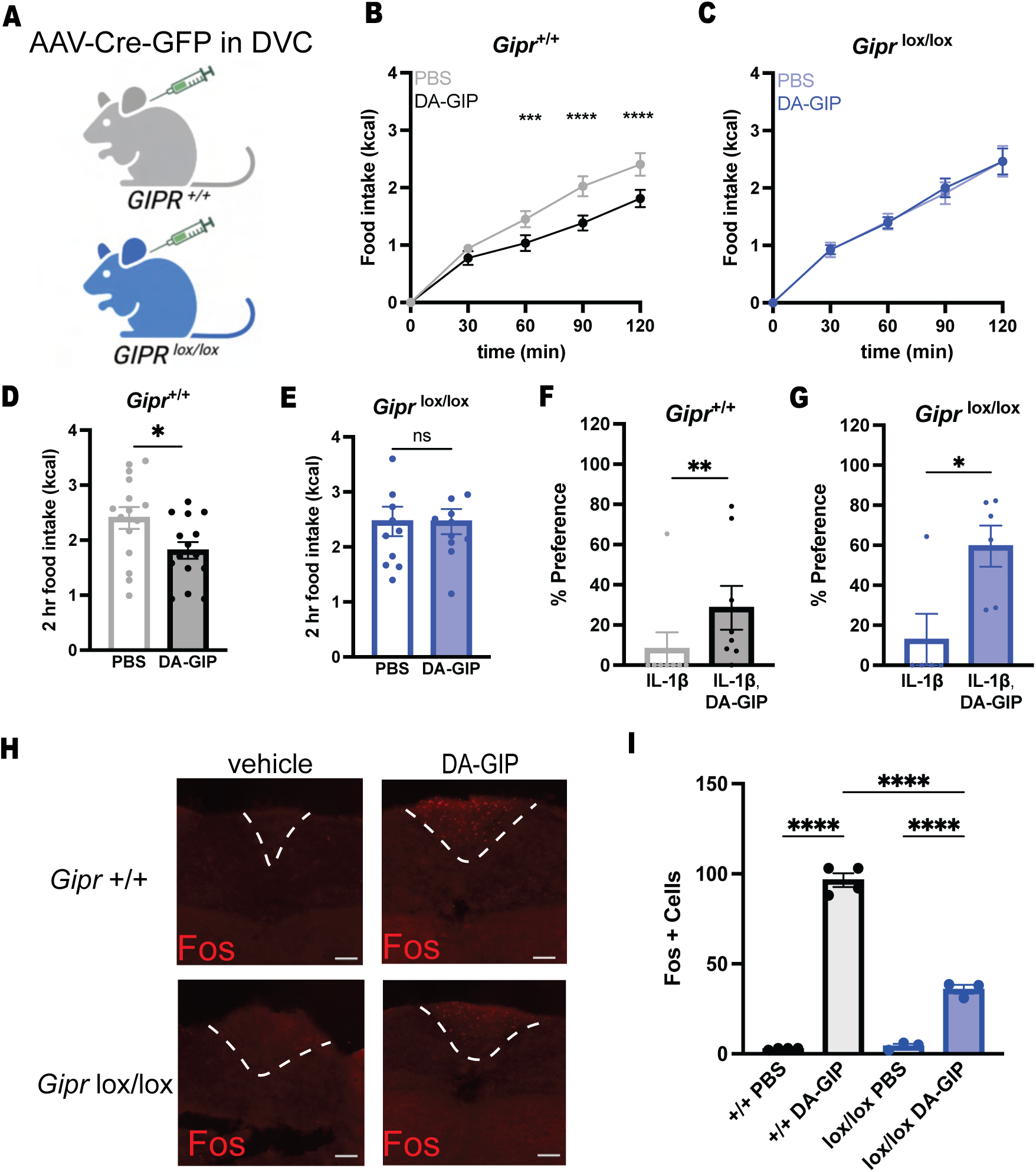
GIPR in DVC are required for DA-GIP-induced anorexia but not anti-aversion. **(A)** Schematic of DVC GIPR knockout approach. **(B, C)** Food intake over time in AAV-Cre-injected *GIPR^+/+^* control mice **(B)** and *GIPR^lox/lox^* mice **(C)** following injection of PBS or DA-GIP (1 mg/kg), i.p. Mice were injected 15 minutes before the start of the dark cycle. Food access commenced at the start of the dark cycle and was measured for 2 hours. N=11-15 mice per group. Two-way ANOVA drug x genotype interaction effect **(B)** p=0.0001. **(C)** p=0.9828. **(D, E)** Two-hour food intake in the mice from **(B, C).** Paired T-test **(D)** p=0.0101; **(E)** p=0.9937. **(F, G)** Percent preference for chocolate versus standard chow in *GIPR^+/+^* control mice **(F)** and *GIPR^lox/lox^*mice **(G)** conditioned with IL-1β (5 μg /kg) or IL-1β + DA-GIP (1 mg/kg) i.p. as indicated. N = 5-8 mice per group. Mann-Whitney (F) p=0.0076; (G) p=0.0216. **(H)** Representative image of Fos staining in the DVC of *GIPR^+/+^* and *GIPR^lox/lox^* mice injected with PBS or DA-GIP i.p. as indicated. Dashed line represents the outline of the AP. Scalebar is 100 µm. **(I)** Quantification of Fos+ cells in the AP of *GIPR^+/+^* and *GIPR^lox/lox^* mice injected with PBS or DA-GIP. N = 3-4 mice per group. Two-way ANOVA drug x genotype interaction effect p<0.0001. **(B, I)** Post-hoc comparisons: ***p<0.001, ****p<0.0001. **(B–G, I)** Data are shown as mean ± SEM.

DVC *Gipr* knockout completely blocked the acute anorectic effect of DA-GIP, indicating that DVC GIPR are necessary for mediating the pharmacologic effects of GIPR agonism on food intake (**Fig. 5B–E**). By contrast, the anti-aversive effect of DA-GIP was intact in both DVC *Gipr* knockout and control mice, suggesting that DVC GIPR are not required for the anti-aversive effect of DA-GIP (**Fig. 5F–G**).

## DISCUSSION

### Distinct neural circuits mediate IL-1β-induced aversion and anorexia

Inflammation induces sickness behaviors, a set of centrally coordinated behavioral changes, including lethargy, loss of appetite (anorexia), and a negative affective state (aversion) [19]. Neurons in the DVC, such as ADCYAP1+ and Tac1+ populations are thought to detect inflammatory and infectious stimuli and coordinate responses such as anorexia and aversion, respectively [23, 26]. DVC neurons drive aversion through projections to the lateral parabrachial nucleus, where CGRP neurons are located [26, 47].

Here we provide evidence that separate neural circuits mediate different aspects of inflammation-induced behavioral responses [26, 44, 46]. Blocking neurotransmission from CGRP neurons completely attenuated IL-1β-induced aversion without rescuing anorexia (**Fig. 1O–P**). By contrast, others have found that silencing CGRP neurons partially rescues anorexia due to lithium chloride, LPS, and GDF-15 [32, 35]. It is possible that silencing CGRP neurons would restore food intake in response to a lower and therefore less anorectic dose of IL-1β. However, here we used the same dose of IL-1β for both CTA and food intake assays, indicating that CGRP neuron activity is required for the development of CTA but not anorexia at a given dose.

We further dissected IL-1β-induced behavioral changes using pharmacologic approaches. We found that the GIPR agonist DA-GIP alleviated behavioral and neural measures of aversion due to IL-1β while enhancing its anorectic effects. OND, when combined with DA-GIP, additively suppressed IL-1β-induced CTA and CGRP neuron activation, but had no effect on food intake. By contrast, the NSAID ketorolac rescued IL-1β-induced anorexia without significantly altering CTA, consistent with prior results [44]. Overall, these findings indicate that GIPR agonism exerts its anti-aversive effects via parabrachial CGRP neurons in a parallel, but possibly convergent pathway to 5-HT_3_ receptor antagonism, and that NSAID treatment blocks the anorectic but not aversive properties of IL-1β. While more work is needed to further unravel the neural circuitry and molecular mechanisms underlying inflammation-induced anorexia and aversion, our findings delineate some key neural circuits driving these processes.

### DVC GIPR are required for GIPR agonist-mediated anorexia but not anti-aversion

GIPR signaling has anti-aversive and anorectic effects, but the underlying mechanisms are incompletely understood [3, 51]. GIPR is expressed in critical homeostatic regions in the brain, including the hypothalamus and DVC [1, 2, 4]. GABAergic neurons are necessary for GIPR agonism-induced reduction in food intake [52], and activation of GIPR-expressing hypothalamic or DVC neuron populations is sufficient to decrease food intake [1, 2]. Chemogenetic activation of GIPR-expressing neurons throughout the DVC induces CTA [1]. By contrast, GIPR-expressing neurons in the AP are necessary for DA-GIP-mediated anti-aversion[13]. Thus, the contributions of DVC GIPR to the behavioral effects of GIP receptor agonism remain unresolved.

Here, we found that selective deletion of DVC GIPR attenuates the anorectic but not the anti-aversive effect of DA-GIP. We also showed that DA-GIP significantly increases Fos expression in the AP, but not the caudal NTS, suggesting that the AP is the primary site of action for DA-GIP in the DVC. Further studies are needed to dissect the contributions of AP versus NTS circuitry in mediating the behavioral and physiologic effects of GIPR agonism, and to determine where GIPR expression is required for anti-aversion. One possibility is that minimal residual GIPR expression in AP neurons in our virus-mediated knockout animal is sufficient to mediate the anti-aversive effect of DA-GIP. Alternatively, GIPR outside the DVC may be responsible for the anti-aversive effect of GIPR signaling. GIPR in GABAergic neurons are required for the anti-aversive effect of DA-GIP [53]; however, GIPR is expressed widely throughout neuronal populations, including the hypothalamus, thalamus, and hippocampus, and peripheral sensory neurons [1, 2, 54]. Future studies will pinpoint where GIPR expression is required for anti-aversion.

### CGRP neuron activation is a robust neural correlate of aversion

CGRP neurons are activated by a wide variety of aversive stimuli, including mechanical, thermal, inflammatory, and pruritic stimuli, as well as cues that predict them [30–35, 55]. Here, we used *in vivo* CGRP neuron activity to substantiate our finding that GIPR agonism alleviates IL-1β-induced aversion. In contrast to DA-GIP, we found that OND did not attenuate IL-1β-induced CTA, consistent with previous studies in rodent models showing that OND did not prevent CTA by serotonin, cisplatin or ipecac [39, 40]. In our study, OND initially decreased IL-1β-induced CGRP neuron activation, but this effect was not significant one hour after injection (**Fig. S3F**). This transience may account for the inability of OND to attenuate CTA, which can form with minute-to hour-long delays between the pairing of a novel flavor and sickness onset [42, 43]. Specifically, CTA is thought to be mediated in part by parabrachial CGRP neurons responding to malaise by activating neurons in the central nucleus of the amygdala that are preferentially activated by novel flavors [56]. OND is a clinically effective anti-nausea and anti-emetic medication [38], suggesting that the CTA assay has limited predictive validity for anti-aversive drugs. By contrast, *in vivo* imaging of CGRP neurons may be a promising screening tool for anti-aversive compounds and may provide a foothold to dissect anti-aversive circuits in future studies.

### Limitations of the study

This study has several limitations. First, an anatomical distinction between AP and NTS-mediated behaviors in response to DA-GIP was not possible using our approach. Fluorescent *in-situ* hybridization demonstrates *Gipr* mRNA expression in the AP and NTS, with stronger *Gipr* probe localization in the AP, and fluorescent-labeled GIPR agonists selectively access circumventricular organs such as the AP over the NTS [1]. Consistent with this, whole-brain Fos analysis in response to short-acting GIPR agonist administration revealed significantly increased Fos expression in the AP, but not the NTS, which is supported by our own findings (**Fig. 5H–I, S6**) [12]. Therefore, it is reasonable to infer that the behavioral effects of DA-GIP at the level of the DVC are mediated predominantly by the AP and not the NTS, though further work is needed to confirm this.

The modest DA-GIP-induced AP Fos expression in mice with GIPR conditionally knocked out of DVC suggests that not all GIPR was deleted in this region and/or that DA-GIP activates some AP neurons indirectly. These possibilities render it impossible to determine for certain that DVC GIPR are dispensable for the anti-aversive effect of DA-GIP (**Fig. 5F, G**). Future studies using transgenic approaches to delete GIPR expression from different subregions of the DVC [57], and from other regions in the brain and periphery, are required to address this question.

Finally, our use of fiber photometry to monitor the activity of CGRP neurons on a population-level may obscure heterogeneity in neural responses to aversive or anti-aversive stimuli within CGRP neurons. Most parabrachial CGRP neurons are uniformly activated by a variety of noxious stimuli [31]; however, it is unclear whether different anti-aversive agents such as OND and DA-GIP impact different subsets of CGRP neurons or have additive effects on the same population of neurons. Future studies using cellular resolution imaging techniques could shed light on this.

## RESOURCE AVAILABILITY

### Lead contact

Further information and requests for resources and reagents should be directed to and will be fulfilled by the lead contact, Lisa Beutler.

### Materials availability

This study did not generate any new unique reagents.

### Data and code availability

- The data reported in the paper will be shared by the lead contact upon request.
- Any additional information required to reanalyze the data reported in this paper is available from the lead contact upon request.
- The code used to analyze fiber photometry data, previously published in [58], has been deposited at Github and is publicly available: https://github.com/nikhayes/fibphoflow.

## ACKNOWLEDGEMENTS

L.R.B. acknowledges support from the American Diabetes Association Pathway to Stop Diabetes Award (12-22-ACE-31) and a McKnight Foundation Neurobiology of Brain Disorders Award. This work was also supported by NIH grants P30-DK020595, K08-DK118188, and R01-DK128477 (L.R.B.) and F30-DK138766 and 5T32-DK007169 (H.S.P.).

## AUTHOR CONTRIBUTIONS

H.S.P. and L.R.B. designed and performed experiments, analyzed data, and prepared the manuscript. N.W.H. produced Python scripts for data analysis. N.W.H., N.L., C.M.L., A.P. and J.L.X., performed experiments.

## METHODS

### Animals

Experimental protocols were approved by the Northwestern University IACUC in accordance with the National Institutes of Health guidelines for the Care and Use of Laboratory Animals (Protocol Numbers: IS00015106, IS00023902, IS00011930, IS00016880, IS00013348, 1S00021524). Mice were housed in a 12/12-h reverse light/dark cycle and given *ad libitum* food and water access unless otherwise stated. Animals were fed *ad libitum* chow (Envigo, 7012, Teklad LM-495 Mouse/Rat Sterilizable Diet). *Calca^tm1.1(cre/EGFP)Rpa^/*J (CGRP-Cre, #033168, Jackson Labs) animals backcrossed onto a C57BL6/J background were used for Tet Tox neural silencing and fiber photometry experiments. *Gipr^lox/lox^* mice were obtained from Dr. Daniel Drucker [50]. C57BL/6 mice were used in food intake and CTA assays where indicated. No statistical methods were used to determine sample sizes. Experiments involved male and female mice 2 to 6 months of age unless otherwise indicated. Male and female mice were used in all experiments and data presented represents both sexes. All experiments were performed during the dark cycle in a dark environment.

### Stereotactic Injections

#### Photometry

For photometry experiments, a recombinant AAV encoding Cre-dependent GCaMP6s (AAV9.CAG.Flex.GCaMP6s, Addgene) was employed as described in [58]. The virus was unilaterally injected into the lateral parabrachial nucleus (LPBN) of CGRP-Cre mice anesthetized with isoflurane at coordinates D/V: –3.81 mm; A/P: −5.18 mm; M/L: +/−1.43 mm from bregma. Some animals were implanted in the left LPBN and others in the right LPBN to ensure responses to stimuli were not lateralized. In addition, a photometry cannula (MFC_400/430– 0.48_3.8mm_MF2.5_FLT, Doric Lenses) was implanted unilaterally in the LPBN at coordinates D/V: –3.71 mm; A/P: −5.18 mm; M/L: +/−1.43 mm from bregma. A bronze mating sleeve (SLEEVE_BR_2.5, Doric Lenses) was also adhered to the implant.

#### Tet Tox

For chronic neuronal silencing experiments, a recombinant AAV encoding Cre-dependent tetanus toxin (AAV9-CMVDIO-eGFP-2A-TeNT) or a control virus (AAV9-CAG-FLEX-eGFP-WPRE) was injected bilaterally in the LPBN of CGRP-Cre mice anesthetized with isoflurane at coordinates D/V: –3.81 mm; A/P: −5.18 mm; M/L: +/−1.43 mm from bregma. Mice without bilateral Tet Tox on histology were excluded from final analysis.

#### DVC Injections

Brainstem injections were based on the protocol described in [59]. GIPR^lox/lox^ mice and GIPR^+/+^ littermate controls were anesthetized via ketamine/xylazine and placed in the stereotaxic frame with the head tilted downwards 90 degrees. The neck muscles were retracted until the brainstem was exposed. The DVC injections used the obex as a reference and injection occurred at coordinates D/V: –0.18 mm; A/P: +0.40 mm; M/L: +/-0.00 mm. 100 nl of AAV9-CMV-HI-CRE-GFP-WPRE was injected at a rate of 1-3 nl/minute.

For all surgeries, mice were treated post-operatively with buprenorphine and meloxicam and kept on a heating pad for observation until they were awake and mobile. Mice were given 2 weeks for viral expression and surgery recovery before experimentation.

### Conditioned taste avoidance (chocolate)

On the days of the conditioning sessions, animals were fasted for 6 hours at the start of the dark cycle (9 am). After fasting, the animals had 30 minutes of habituation in individual cages, then were given chocolate for 30 minutes and their intake was measured. At the end of the 30 min, animals received i.p. injection of IL-1β or vehicle with or without an anti-aversive agent and were returned to their home cages. The mice were allowed one recovery day before a second identical conditioning session, and two recovery days between the final conditioning session and the preference testing session. On the day of the preference test, the animals underwent a 6 hour fast followed by 30-minute access to their standard chow and chocolate. The intake was recorded in grams and converted to kcal and the percent preference for chocolate was calculated as (chocolate kcal / total kcal consumed * 100%).

### Conditioned taste avoidance (saccharin)

Group-housed mice were acclimated to 2 water bottles for 7 days in their home cages. Then, mice were water-deprived for 16 hours (overnight). On days 1 and 2, mice had access to water for 30 minutes in individual cages at the start of the dark cycle and for 2 hours in the afternoon, prior to being water-restricted again. On day 3, mice were given 30-minute access to a novel 0.15% saccharin solution, followed by an injection of PBS or IL-1β. Water was available for 1 h in the afternoon on day 3 and again on days 4 and 6, following the same schedule as days 1 and 2. Conditioning was repeated on day 5. On day 7, mice were given access to two bottles (water and 0.15% saccharin) for 30 min. The volume consumed was recorded in grams and converted to milliliters and the percent preference for saccharin was calculated.

### Food intake experiments

Mice were habituated to feeding cages, handling, and intraperitoneal injections for 1 week before food intake experiments. The day before the experiment, mice were fasted for 16 hours (overnight) then allowed to acclimate in individual feeding chambers for approximately 30 minutes. Mice were then injected with the following compounds and doses as indicated in the text and figures: IL-1β (5 μg/kg, Biolegend 575106), DA-GIP (1 mg/kg, Bachem 4054476), ondansetron (1 mg/kg, Tocris 99614-01-4), and ketorolac (15 mg/kg, Sigma-Aldrich 74103-07-4). IL-1β was injected at the same time as DA-GIP or ondansetron. Ketorolac was injected 30 minutes prior to IL-1β or PBS based on prior studies [44, 46]. Mice were re-fed after injections were completed. For C57BL6/J studies, food intake was monitored at 15, 30, 45, 60, 90, and 120 min after food presentation. For Tet Tox studies, food intake was monitored at 1, 2, 4, and 6 hours after food presentation. The order of drug condition the mice were exposed to was always counterbalanced across mice.

### CGRP neuron photometry

Fiber photometry was performed as we have described previously [48] [58]. Two different photometry processors were used for data collection in this study. One photometry rig includes LEDs and LED driver separate from the processor (RZ5P, TDT (processor); DC4100 (LED driver); M405FP1 and M470F3 (LEDs), Thorlabs). The second rig has these components integrated into the processor (RZ10X, TDT). The neural activity of each mouse was recorded using the same system and patch cord for every session allowing for reliable within-mouse comparisons of calcium signal over time. Continuous blue LED (465–470 nm) and UV LED (405 nm) served as excitation light sources. These LEDs were modulated at distinct rates and delivered to a filtered minicube (Doric Lenses) before connecting through patch cords (MFP_400/430/1100– 0.57_2m_FCM-MF2.5_LAF, Doric Lenses) to mouse implants (MFC_400/430– 0.48_3.8mm_MF2.5_FLT, Doric Lenses). GCaMP6s calcium signals and isosbestic signals were collected through the same fibers back to dichroic ports of a minicube and transmitted to photoreceivers (Newport Visible Femtowatt Photoreceiver for the RZ5P system; integrated Lux photosensors in the RZ10X system). Digital signals sampled at 1.0173 kHz were then demodulated, lock-in amplified, and collected through the processor (RZ5P or RZ10X, TDT). Data were collected using the software Synapse (TDT), exported, and analyzed using custom Python code and Prism software.

Animals were habituated to the recording chambers for 15 minutes during fiber photometry recording before intraperitoneal injections. Photometry recording continued for 60 minutes following the injection. Unless otherwise specified, mice were fasted overnight prior to recordings.

CGRP neurons are less active in fasted mice than sated; therefore, fasted animals are more suited to studying increases in CGRP neuron activity [31] [55]. Mice that did not exhibit a minimal 15% increase in CGRP neuron calcium activity in response to toe pinch were considered technical failures and excluded from future experiments based on previously published work showing that CGRP neurons and/or glutamatergic LPBN neurons are responsive to toe pinch [31, 60]. For experiments assessing CGRP neuron response to inflammatory and anti-aversive agents, the order of drug and vehicle treatment was counterbalanced across mice. Drugs were administered at the following doses: IL-1β (10 μg/kg), DA-GIP (1 mg/kg), OND (1 mg/kg), KET (15 mg/kg), LPS (0.5 mg/kg, Sigma-Aldrich 93572-42-0), and TNF-α (0.2 mg/kg, BioLegend 575208).

### DVC Fos expression

For Fos studies, *Gipr*^lox/lox^ and *Gipr*^+/+^ mice that underwent AAV-Cre-GFP delivery into the DVC at least two weeks earlier were injected with PBS vehicle control or DA-GIP i.p. After 1 hour, brains were dissected and underwent staining for Fos as described below. Only animals with at least 1-2 intact sections of the DVC containing the AP and surrounding bilateral NTS were included in the analysis. Images were obtained at 10x using consistent settings across samples on the Leica Thunder Imaging System and then imported into ImageJ, where Fos positive cells were counted. For animals with multiple sections containing the AP and surrounding NTS, the count was averaged across the sections. The experimenter was blinded to treatment condition until cell counting was complete.

### Immunohistochemistry

Immunohistochemistry was performed as we have described previously [58]. To confirm virus-induced Cre-GFP expression in the dorsal vagal complex, brains were dissected, postfixed for 12-48 hours in 4% paraformaldehyde, then placed in 30% sucrose for one or two days for cryoprotection. Tissues were frozen in Optimal Cutting Temperature embedding compound (4585, Fisher HealthCare) at −30°C until sectioned. Free-floating sections (40 μm) were made using a cryostat. The sections were washed, blocked (10% normal donkey serum (017-000-121, Jackson ImmunoResearch) and 0.3% Triton X-100 (TX1568-1, EMD Millipore) in PBS) for 1 h at room temperature, and then incubated with a primary antibody (rabbit anti-GFP, Invitrogen, A11122, 1:500 or mouse anti-GFP, Cell Signaling Technology, 2955S, 1:500) overnight at 4°C. Samples were then washed and incubated with a secondary antibody (donkey anti-rabbit Alexa Fluor 488; Jackson ImmunoResearch, 711-545-152, 1:500 or donkey anti-mouse Alexa Fluor 488; Jackson ImmunoResearch, 715-545-150, 1:500) for 2 h at room temperature before being mounted and imaged on a Leica Thunder Imaging System.

To stain for Fos, animals were injected with PBS vehicle control or DA-GIP i.p. After 1 hour, brains were dissected and processed and imaged as described above. The sections were washed, blocked (10% normal donkey serum (017-000-121, Jackson ImmunoResearch) and 0.3% Triton X-100 (TX1568-1, EMD Millipore) in PBS) for 1 h at room temperature, and then incubated with a primary antibody (rabbit anti-c-Fos, Cell Signaling Technology, 2250, 1:500) overnight at 4°C. Samples were then washed and incubated with a secondary antibody (donkey anti-rabbit Alexa Fluor 594; Jackson ImmunoResearch, 711-585-152, 1:500) for 2 h at room temperature.

### Statistical Analysis

#### Photometry

Normalization of CGRP neuron activity post-injection compared to baseline activity was performed on calcium-dependent emission and isosbestic control signals via the formula: ΔF/F = (Ft – F0)/F0, where Ft represents fluorescence at time (t), and F0 represents the average fluorescence during the 5-min baseline period preceding time zero (stimulus start time). ΔF/F (%) refers the mean ΔF_t_/F_0∗_100. Bar graphs quantifying CGRP neural responses to i.p. injection represent the average ΔF/F (%) between 50-60 minutes after injection (**Figure 1, 1S**) or over a 1-min period 15, 30, 45, and 60 minutes after injection (**Figures 2, 3, S3, S4**).

#### Conditioned taste avoidance in C57BL6/J mice

One-way ANOVA with post-hoc Holm-Šidák was used to assess differences in preference across drug treatments in CTA assays (**Figures 2, 3, S3, S4**).

#### Conditioned taste avoidance in tetanus toxin mice

Two-way ANOVA with post-hoc Holm-Šidák was used to assess differences in preference between different virus and drug treatment conditions in the CTA assay (**Figures 1, S1**).

#### Conditioned taste avoidance in *Gipr*^lox/lox^ and *Gipr*^+/+^ mice

The non-parametric Mann-Whitney test was used to assess differences in preference within genotype across different drug treatment conditions in CTA assays (**Figure 5**).

#### Food intake in C57BL6/J mice

One-way repeated-measures ANOVA was used to assess differences in food intake between the treatment conditions (**Figure 4**).

#### Food intake in tetanus toxin mice

Two-way ANOVA was used to assess differences in food intake between different virus and treatment conditions (**Figure 1**).

#### Fos + cell count in AP of *Gipr*^lox/lox^ and *Gipr*^+/+^ mice

Two-way ANOVA with post-hoc Holm-Šidák testing was used to assess differences in Fos expression across genotypes and treatment conditions (**Figure 5**).

#### Fos + cell count in NTS of *Gipr*^+/+^ mice

Unpaired T-test was used to assess the difference in Fos expression between drug conditions (**Figure S6**).

Prism was used for all statistical analyses, and significance was defined as p < 0.05. Sample sizes are indicated in the figure legends for each experiment. The Holm-Šídák multiple comparisons test was used in conjunction with ANOVA where appropriate. Where multiple technical replicates of an experiment were performed, trials from the same animal were averaged and handled as a single biological replicate for data analysis and visualization.

## FIGURE LEGENDS

**Supplemental Figure 1.**
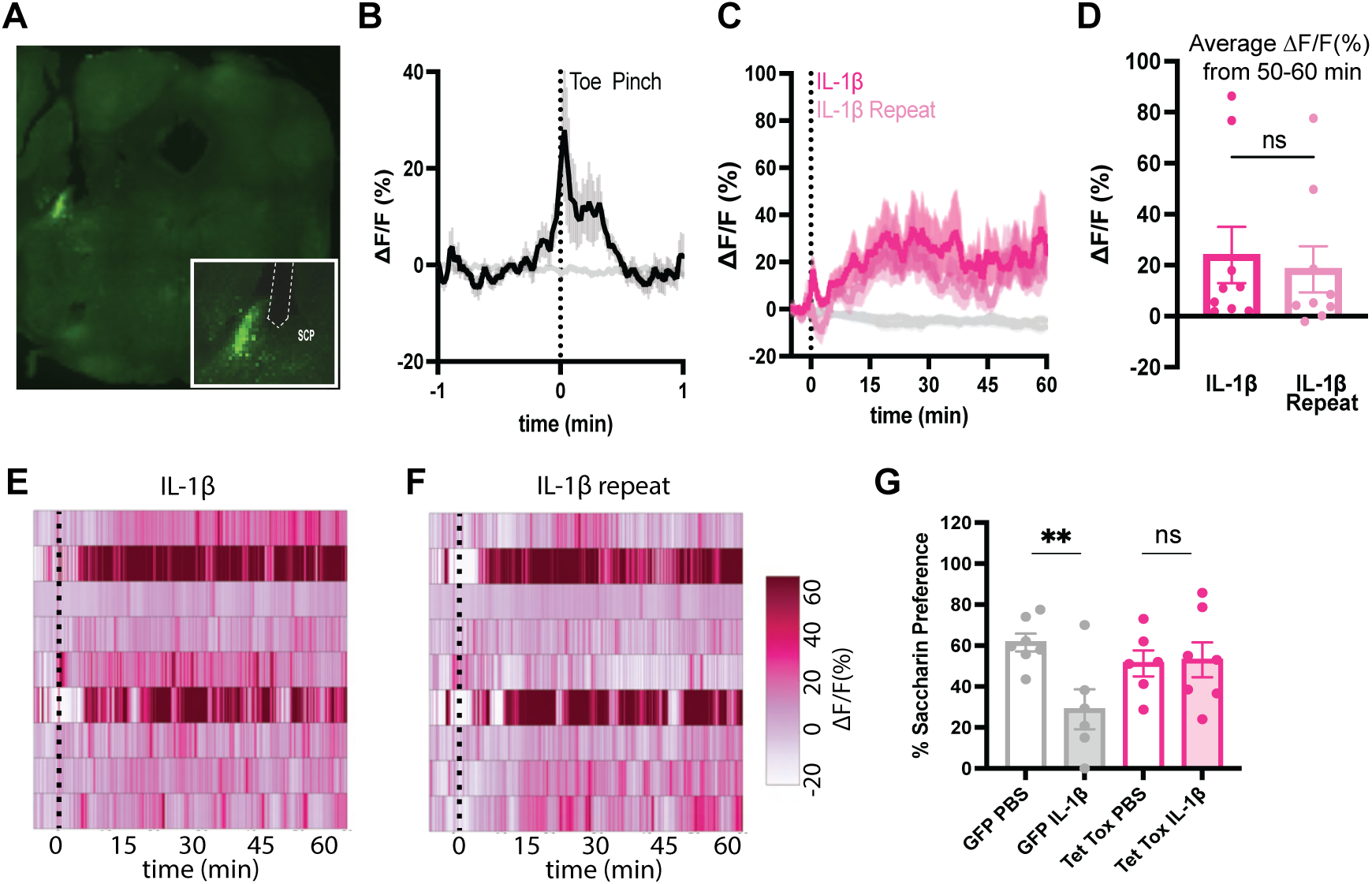
Histological and functional validation of fiber photometry in CGRP neurons. (**A**) Coronal section from a *Calca^Cre:GFP^* mouse depicting GCaMP expression (green) and path of optical fiber in inset. SCP = superior cerebellar peduncle. Inset scale bar represents 100 μm. Dashed white line indicates implant insertion site. (**B**) Calcium signal in CGRP neurons from mice undergoing a toe pinch. N = 9 mice. (**C**) Calcium signal in CGRP neurons from fasted mice injected with IL-1β (10 µg/kg) i.p. on two separate occasions between 3 and 14 days apart. N = 9 mice per group. (**D**) Average ΔF/F in mice from **(C)** 50-60 minutes after injection. Paired T test. p=0.2225. **(E–F)** Heat maps showing ΔF/F in individual mice injected with first and repeated IL-1β. Each heatmap line represents an individual animal. **(G)** Percent preference for saccharin-sweetened versus standard water in GFP and TetTox-injected *Calca^Cre:GFP^* mice conditioned with PBS or IL-1β (5 μg /kg) i.p. as indicated. N = 6-7 mice per group. Two-way ANOVA drug x virus interaction p=.0312. Post-hoc comparisons: *p<0.05, **p<0.01. **(B, C)** Vertical dashed lines indicate the time of toe pinch or injection. Traces indicate mean ± SEM. **(D, G)** Data are shown as mean ± SEM.

**Supplemental Figure 2.**
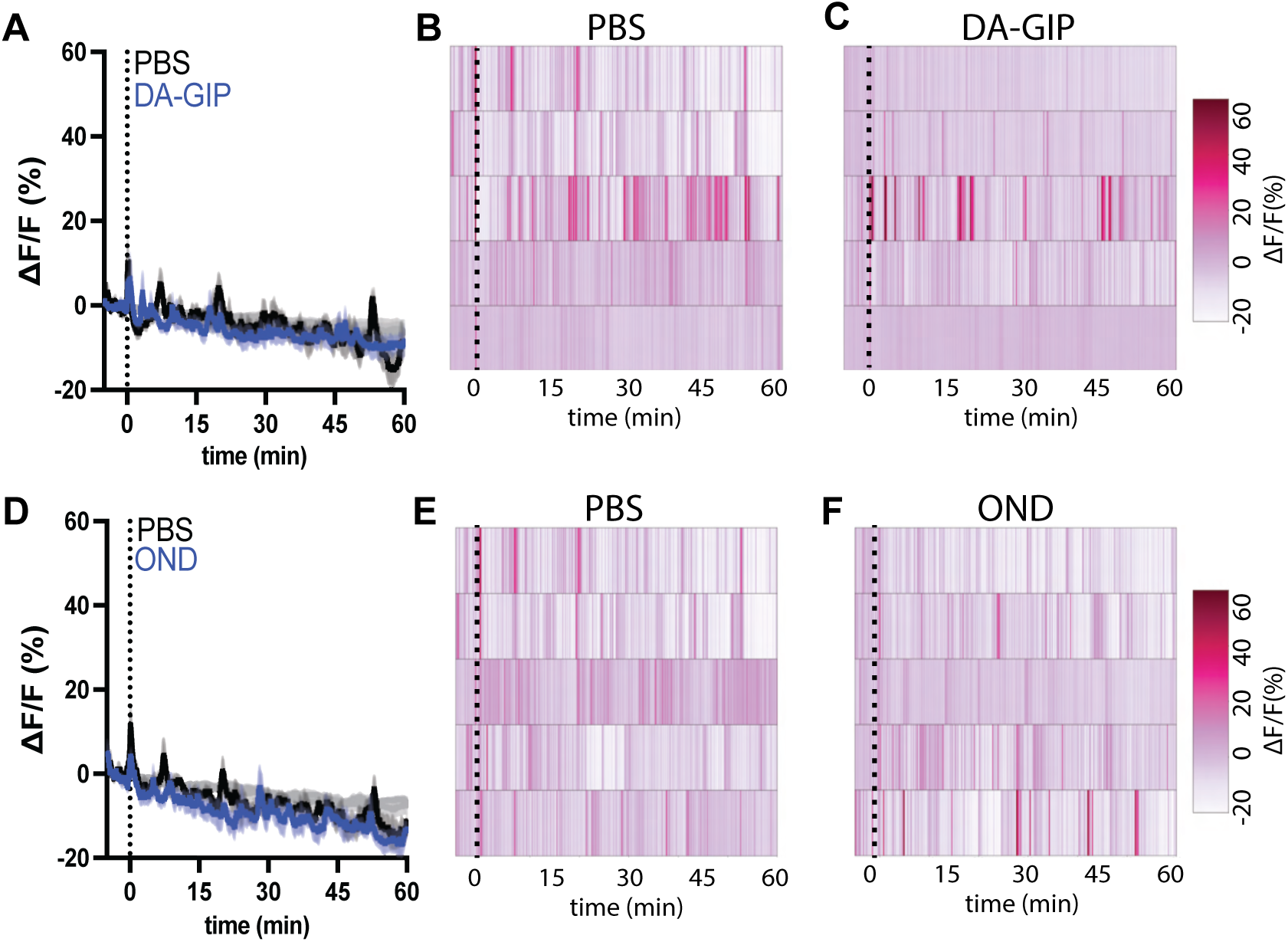
Neither DA-GIP nor ondansetron impact CGRP neuron activity in fed mice. **(A, D)** Calcium signal in CGRP neurons from fed mice injected with PBS (black traces) or DA-GIP (1 mg/kg, **A**), or OND (1 mg/kg, **D**) i.p. N = 5 mice per group. Two-way ANOVA effect of drug **(A)** p=0.5; **(D)** p=0.13. Isosbestic traces for all recordings are shown in gray. Vertical dashed lines indicate the time of injection. Traces indicate mean ± SEM. **(B, C, E, F)** Heat maps showing ΔF/F in individual mice from **(A, D)** injected with PBS, DA-GIP, or OND as indicated. Each heatmap line represents an individual animal.

**Supplemental Figure 3.**
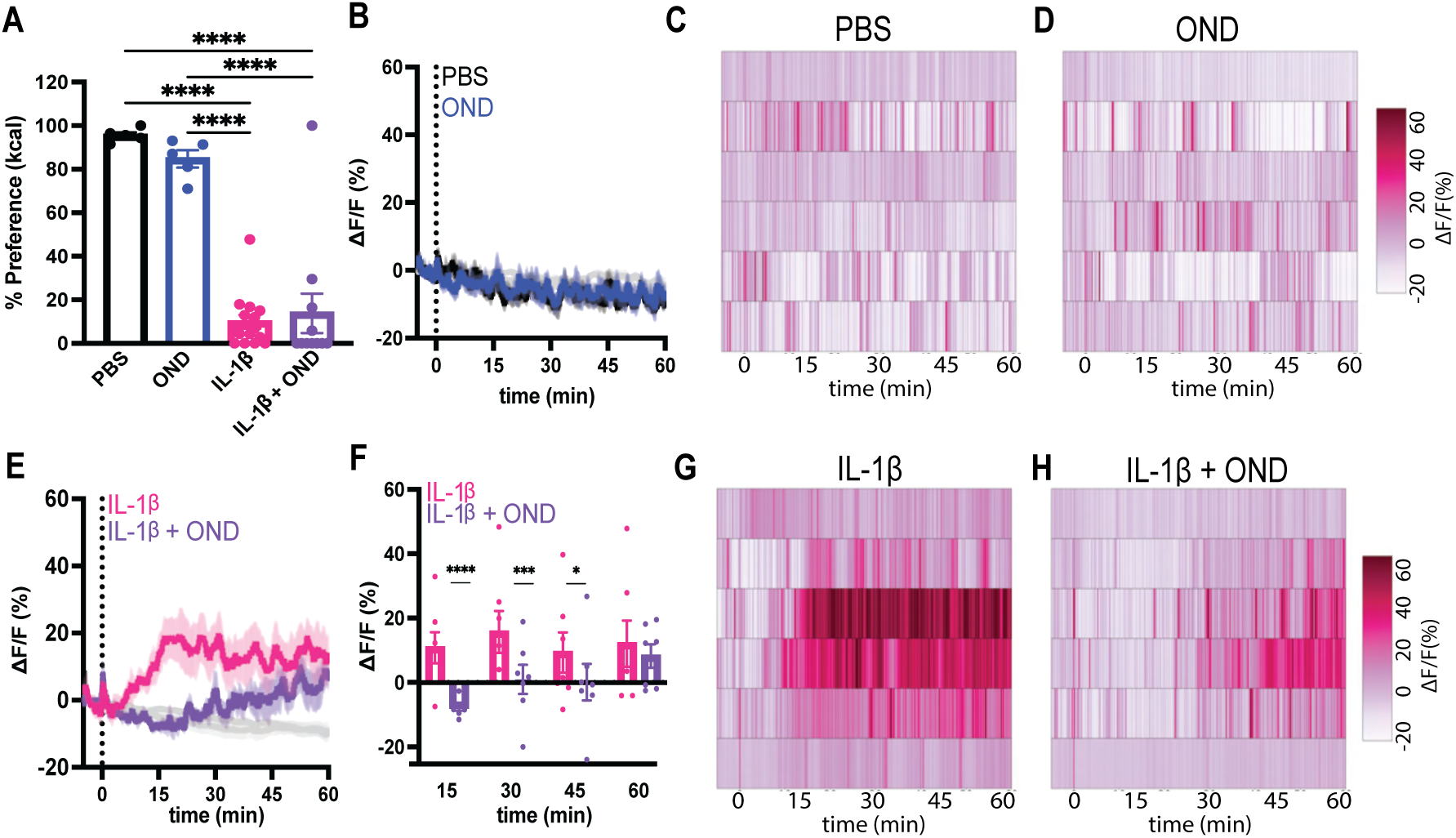
Ondansetron decreases IL-1β-induced CGRP neuron activation without affecting CTA. **(A)** Percent preference for chocolate versus standard chow in C57BL/6 mice conditioned with PBS, OND (1 mg/kg), IL-1β (5 μg /kg), or IL-1β + OND i.p. as indicated N = 5-15 mice per group. One-way ANOVA p< 0.0001. **(B)** Calcium signal in CGRP neurons from fasted mice injected with PBS or OND (1 mg/kg) i.p. N = 6 mice per group. **(C, D)** Heat maps showing ΔF/F in individual mice from **(B)** injected with PBS or OND as indicated. (**E**) Calcium signal in CGRP neurons from fasted mice injected with IL-1β (10 µg/kg) or IL-1β + OND (1 mg/kg) i.p. N = 7 mice per group. (F) Average ΔF/F in mice from **(E)** 15, 30, 45, and 60 minutes after injection. Two-way ANOVA effect of drug p=0.0392. **(G, H)** Heat maps showing ΔF/F in individual mice injected with IL-1β or IL-1β + OND as indicated. **(A, F)** Data are shown as mean ± SEM. Post-hoc comparisons: *p<0.05, **p < 0.01; ***p<0.001, ****p<0.0001. **(B, E)** Isosbestic traces for all recordings are shown in gray. Vertical dashed lines indicate the time of injection. Traces indicate mean ± SEM. **(C, D, G, H)** Each heatmap line represents an individual animal.

**Supplemental Figure 4.**
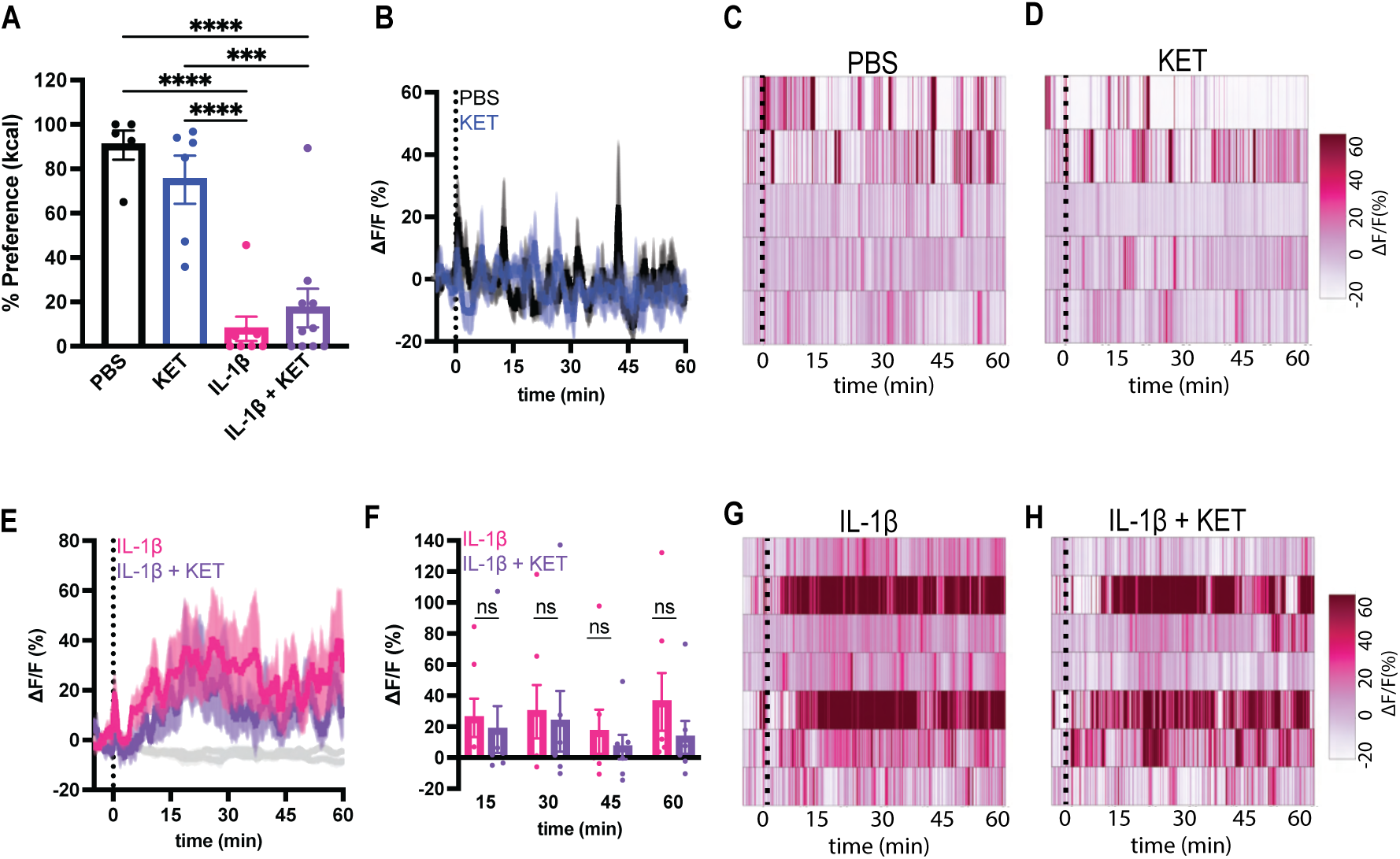
Ketorolac does not decrease IL-1β-induced CGRP neuron activation or CTA. **(A)** Percent preference for chocolate versus standard chow in C57BL/6 mice conditioned with PBS, KET (15 mg/kg), IL-1β (5 μg /kg), or IL-1β + KET i.p as indicated. N = 5-10 mice per group. One-way ANOVA p< 0.0001. Post-hoc comparisons: ***p<0.001, ****p<0.0001. **(B)** Calcium signal in CGRP neurons from fasted mice injected with PBS or KET (15 mg/kg) i.p. N = 5 mice per group. **(C–D)** Heat maps showing ΔF/F in individual mice injected with PBS or KET. **(E)** Calcium signal in CGRP neurons from fasted mice injected with IL-1β (10 µg/kg) or IL-1β after pre-treatment with KET (15 mg/kg) i.p. N = 7 mice per group. **(F)** Average ΔF/F in mice from **(E)** 15, 30, 45, and 60 minutes after injection. Two-way ANOVA effect of drug p = 0.3964. **(G–H)** Heat maps showing ΔF/F in individual mice injected with IL-1β or IL-1β + KET. **(A, F)** Data are shown as mean ± SEM. **(B, E)** Isosbestic traces for all recordings are shown in gray. Vertical dashed lines indicate the time of injection. Traces indicate mean ± SEM. **(C, D, G, H)** Each heatmap line represents an individual animal.

**Supplemental Figure 5.**
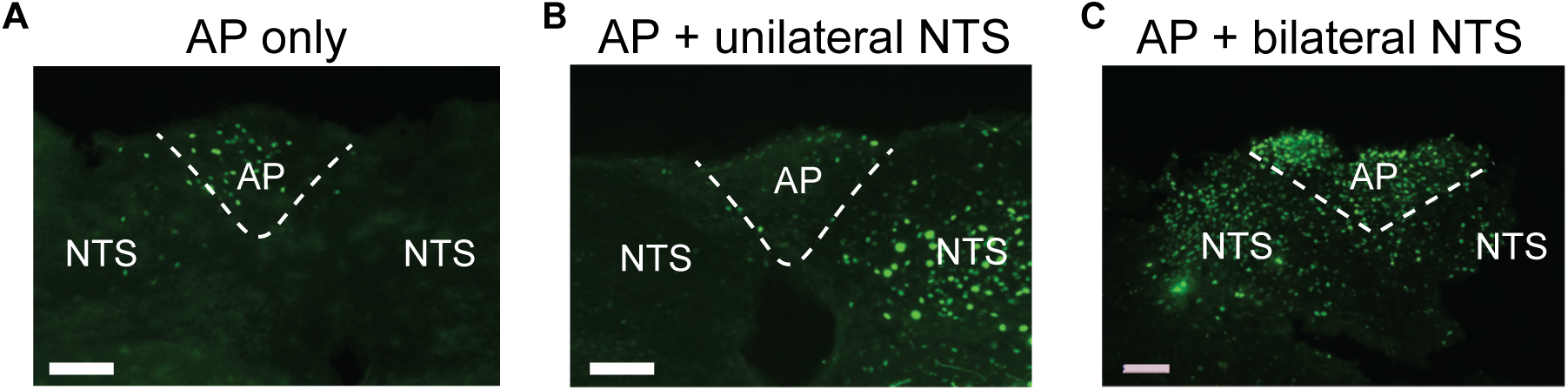
Patterns of AAV9-Cre-GFP expression across GIPR^lox/lox^ DVC. **(A)** Representative image of Cre-GFP expression primarily in AP neurons in a *GIPR^lox/lox^* mouse. **(B)** Representative image of Cre-GFP expression in AP and unilateral NTS neurons in a *GIPR^lox/lox^* mouse. **(C)** Representative image of Cre-GFP expression in AP and bilateral NTS neurons in a *GIPR^lox/lox^* mouse. **(A–C)** Scale bar is 100 µm.

**Supplemental Figure 6.**
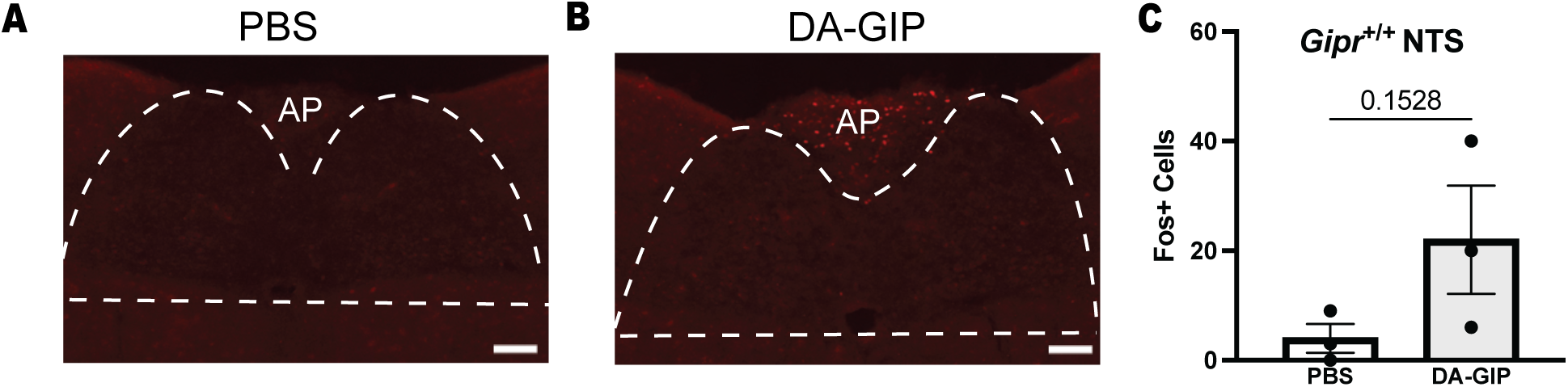
DA-GIP does not significantly increase Fos expression in caudal NTS at the level of the AP. **(A)** Representative image of Fos staining in the DVC of a *GIPR^+/+^* mouse injected with PBS i.p. **(B)** Representative image of Fos staining in the DVC of a *GIPR^+/+^* mouse injected with DA-GIP i.p. **(C)** Quantification of Fos+ cells in the NTS of *GIPR^+/+^* mice injected with PBS or DA-GIP. N = 3 mice per group. Unpaired T-test p=0.1528. Data are shown as mean ± SEM. **(A, B)** Dashed line represents outline of NTS. Scalebar is 100 µm.

## REFERENCES

1. Adriaenssens, A., et al., Hypothalamic and brainstem glucose-dependent insulinotropic polypeptide receptor neurons employ distinct mechanisms to a6ect feeding. JCI Insight, 2023. 8(10).

2. Adriaenssens, A.E., et al., Glucose-Dependent Insulinotropic Polypeptide Receptor-Expressing Cells in the Hypothalamus Regulate Food Intake. Cell Metab, 2019. 30(5): p. 987–996 e6.

3. Campbell, J.E., Targeting the GIPR for obesity: To agonize or antagonize? Potential mechanisms. Mol Metab, 2021. 46: p. 101139.

4. Zhang, Q., et al., The glucose-dependent insulinotropic polypeptide (GIP) regulates body weight and food intake via CNS-GIPR signaling. Cell Metab, 2021. 33(4): p. 833–844 e5.

5. Gutgesell, R.M., et al., GIPR agonism and antagonism decrease body weight and food intake via di6erent mechanisms in male mice. Nat Metab, 2025.

6. JastreboM, A.M., et al., Tirzepatide Once Weekly for the Treatment of Obesity. N Engl J Med, 2022. 387(3): p. 205–216.

7. Killion, E.A., et al., Anti-obesity e6ects of GIPR antagonists alone and in combination with GLP-1R agonists in preclinical models. Sci Transl Med, 2018. 10(472).

8. Mroz, P.A., et al., Optimized GIP analogs promote body weight lowering in mice through GIPR agonism not antagonism. Mol Metab, 2019. 20: p. 51–62.

9. Svendsen, B., et al., Pharmacological antagonism of the incretin system protects against diet-induced obesity. Mol Metab, 2020. 32: p. 44–55.

10. Veniant, M.M., et al., A GIPR antagonist conjugated to GLP-1 analogues promotes weight loss with improved metabolic parameters in preclinical and phase 1 settings. Nat Metab, 2024. 6(2): p. 290–303.

11. Costa, A., et al., Anorectic and aversive e6ects of GLP-1 receptor agonism are mediated by brainstem cholecystokinin neurons, and modulated by GIP receptor activation. Mol Metab, 2022. 55: p. 101407.

12. Samms, R.J., et al., GIPR Agonism Inhibits PYY-Induced Nausea-Like Behavior. Diabetes, 2022. 71(7): p. 1410–1423.

13. Zhang, C., et al., A brainstem circuit for nausea suppression. Cell Rep, 2022. 39(11): p. 110953.

14. Borner, T., et al., GIP Receptor Agonism Attenuates GLP-1 Receptor Agonist-Induced Nausea and Emesis in Preclinical Models. Diabetes, 2021. 70(11): p. 2545–2553.

15. Borner, T., et al., GIP receptor agonism blocks chemotherapy-induced nausea and vomiting. Mol Metab, 2023. 73: p. 101743.

16. Knop, F.K., et al., A long-acting glucose-dependent insulinotropic polypeptide receptor agonist improves the gastrointestinal tolerability of glucagon-like peptide-1 receptor agonist therapy. Diabetes Obes Metab, 2024. 26(11): p. 5474–5478.

17. Morimoto, T.A.N.N.K.A.T., Gip receptor activating peptide. 2018.

18. Dantzer, R., Evolutionary Aspects of Infections: Inflammation and Sickness Behaviors. Curr Top Behav Neurosci, 2023. 61: p. 1–14.

19. Hart, B.L., Biological basis of the behavior of sick animals. Neurosci Biobehav Rev, 1988. 12(2): p. 123–37.

20. Wang, A., et al., Opposing E6ects of Fasting Metabolism on Tissue Tolerance in Bacterial and Viral Inflammation. Cell, 2016. 166(6): p. 1512–1525 e12.

21. Balestrieri, P., et al., Nutritional Aspects in Inflammatory Bowel Diseases. Nutrients, 2020. 12(2).

22. Dantzer, R., et al., From inflammation to sickness and depression: when the immune system subjugates the brain. Nat Rev Neurosci, 2008. 9(1): p. 46–56.

23. Ilanges, A., et al., Brainstem ADCYAP1(+) neurons control multiple aspects of sickness behaviour. Nature, 2022. 609(7928): p. 761–771.

24. Jin, H., et al., A body-brain circuit that regulates body inflammatory responses. Nature, 2024. 630(8017): p. 695–703.

25. Osterhout, J.A., et al., A preoptic neuronal population controls fever and appetite during sickness. Nature, 2022. 606(7916): p. 937–944.

26. Xie, Z., et al., The gut-to-brain axis for toxin-induced defensive responses. Cell, 2022. 185(23): p. 4298–4316 e21.

27. Zhao, R., H. Zhou, and S.B. Su, A critical role for interleukin-1beta in the progression of autoimmune diseases. Int Immunopharmacol, 2013. 17(3): p. 658–69.

28. Hellerstein, M.K., et al., Interleukin-1-induced anorexia in the rat. Influence of prostaglandins. J Clin Invest, 1989. 84(1): p. 228–35.

29. Plata-Salaman, C.R., Meal patterns in response to the intracerebroventricular administration of interleukin-1 beta in rats. Physiol Behav, 1994. 55(4): p. 727–33.

30. Campos, C.A., et al., Cancer-induced anorexia and malaise are mediated by CGRP neurons in the parabrachial nucleus. Nat Neurosci, 2017. 20(7): p. 934–942.

31. Campos, C.A., et al., Encoding of danger by parabrachial CGRP neurons. Nature, 2018. 555(7698): p. 617–622.

32. Carter, M.E., et al., Genetic identification of a neural circuit that suppresses appetite. Nature, 2013. 503(7474): p. 111–4.

33. Chen, J.Y., et al., Parabrachial CGRP Neurons Establish and Sustain Aversive Taste Memories. Neuron, 2018. 100(4): p. 891–899 e5.

34. Jagot, F., et al., The parabrachial nucleus elicits a vigorous corticosterone feedback response to the pro-inflammatory cytokine IL-1beta. Neuron, 2023. 111(15): p. 2367–2382 e6.

35. Sabatini, P.V., et al., GFRAL-expressing neurons suppress food intake via aversive pathways. Proc Natl Acad Sci U S A, 2021. 118(8).

36. Goehler, L.E., et al., Blockade of cytokine induced conditioned taste aversion by subdiaphragmatic vagotomy: further evidence for vagal mediation of immune-brain communication. Neurosci Lett, 1995. 185(3): p. 163–6.

37. Tazi, A., et al., Interleukin-1 induces conditioned taste aversion in rats: a possible explanation for its pituitary-adrenal stimulating activity. Brain Res, 1988. 473(2): p. 369–71.

38. Cubeddu, L.X., et al., E6icacy of ondansetron (GR 38032F) and the role of serotonin in cisplatin-induced nausea and vomiting. N Engl J Med, 1990. 322(12): p. 810–6.

39. Mele, P.C., et al., Cisplatin-induced conditioned taste aversion: attenuation by dexamethasone but not zacopride or GR38032F. Eur J Pharmacol, 1992. 218(2-3): p. 229–36.

40. Rudd, J.A., M.P. Ngan, and M.K. Wai, 5-HT3 receptors are not involved in conditioned taste aversions induced by 5-hydroxytryptamine, ipecacuanha or cisplatin. Eur J Pharmacol, 1998. 352(2-3): p. 143–9.

41. McAllister, K.H. and J.A. Pratt, GR205171 blocks apomorphine and amphetamine-induced conditioned taste aversions. Eur J Pharmacol, 1998. 353(2-3): p. 141–8.

42. Garcia, J.E. F.R.; Koelling, R.A., Learning with prolonged delay of reinforcement. Psychonomic Science, 1966. 5(3): p. 121–122.

43. Revusky, S.H. and E.W. Bedarf, Association of illness with prior ingestion of novel foods. Science, 1967. 155(3759): p. 219–20.

44. Nilsson, A., et al., Inflammation-induced anorexia and fever are elicited by distinct prostaglandin dependent mechanisms, whereas conditioned taste aversion is prostaglandin independent. Brain Behav Immun, 2017. 61: p. 236–243.

45. Uehara, A., et al., Indomethacin blocks the anorexic action of interleukin-1. Eur J Pharmacol, 1989. 170(3): p. 257–60.

46. Cheng, W., et al., NTS Prlh overcomes orexigenic stimuli and ameliorates dietary and genetic forms of obesity. Nat Commun, 2021. 12(1): p. 5175.

47. Fritz, M., et al., Prostaglandin-dependent modulation of dopaminergic neurotransmission elicits inflammation-induced aversion in mice. J Clin Invest, 2016. 126(2): p. 695–705.

48. Huang, K.P., et al., Dissociable hindbrain GLP1R circuits for satiety and aversion. Nature, 2024.

49. McMorrow, H.E., et al., Incretin hormones and pharmacomimetics rapidly inhibit AgRP neuron activity to suppress appetite. bioRxiv, 2024.

50. Zhang, C., et al., Area Postrema Cell Types that Mediate Nausea-Associated Behaviors. Neuron, 2021. 109(3): p. 461–472 e5.

51. Campbell, J.E., et al., TCF1 links GIPR signaling to the control of beta cell function and survival. Nat Med, 2016. 22(1): p. 84–90.

52. Borner, T., B.C. De Jonghe, and M.R. Hayes, The antiemetic actions of GIP receptor agonism. Am J Physiol Endocrinol Metab, 2024. 326(4): p. E528–E536.

53. Liskiewicz, A., et al., Glucose-dependent insulinotropic polypeptide regulates body weight and food intake via GABAergic neurons in mice. Nat Metab, 2023. 5(12): p. 2075–2085.

54. Wean, J., et al., Specific loss of GIPR signaling in GABAergic neurons enhances GLP-1R agonist-induced body weight loss. Mol Metab, 2025. 95: p. 102074.

55. Okawa, T., et al., Sensory and motor physiological functions are impaired in gastric inhibitory polypeptide receptor-deficient mice. J Diabetes Investig, 2014. 5(1): p. 31–7.

56. Campos, C.A., et al., Parabrachial CGRP Neurons Control Meal Termination. Cell Metab, 2016. 23(5): p. 811–20.

57. Zimmerman, C.A., et al., A neural mechanism for learning from delayed postingestive feedback. Nature, 2025.

58. Yacawych, W.T., et al., A single dorsal vagal complex circuit mediates the aversive and anorectic responses to GLP1R agonists. bioRxiv, 2025.

59. Lorch, C.M., et al., Sucrose overconsumption impairs AgRP neuron dynamics and promotes palatable food intake. Cell Rep, 2024. 43(2): p. 113675.

60. Joshi, K., et al., Stereotaxic Surgical Approach to Microinject the Caudal Brainstem and Upper Cervical Spinal Cord via the Cisterna Magna in Mice. J Vis Exp, 2022(179).

61. Sun, L., et al., Parabrachial nucleus circuit governs neuropathic pain-like behavior. Nat Commun, 2020. 11(1): p. 5974.

